# Sublingual Buprenorphine/Naloxone Film in Unanesthetized Rats: Feasibility, Tolerability, and Plasma Exposure

**DOI:** 10.64898/2026.10.30.763075

**Authors:** Yao Wang, Qiang Chen, Thomas Deakin, Trent Emerick, Yan Xu, Pei Tang

**Author notes:** Corresponding Author: Pei Tang, Corresponding Author Current Address: 2049 Biomedical Science Tower 3, University of Pittsburgh, Pittsburgh PA, 15261 USA.

## Abstract

**Background:** Buprenorphine/naloxone sublingual film is first-line treatment for opioid use disorder, yet preclinical rodent studies use routes that do not match clinical dosing. We tested sublingual film administration in unanesthetized rats.

**Methods:** Male Sprague-Dawley rats received film delivering 3.2 mg/kg buprenorphine, scaled from the 16 mg/day clinical dose by body surface area and corrected for species differences in sublingual bioavailability. Plasma buprenorphine was measured by LC-MS/MS after a single dose. Body weight, liver weight, serum AST, ALT and ALP, and liver histopathology were compared with controls after 21-days of once-daily dosing.

**Results:** Plasma buprenorphine peaked at 4.0 ng/mL at 2 h and was 0.9 ng/mL at 24 h, with an AUC_0-24_ of 39.0 h·ng/mL, spanning concentrations reported as therapeutic in patients. Repeat dosing reduced absolute and relative liver weight and relative liver weight, with no change in serum enzymes or liver histology.

**Conclusions:** Sublingual film dosing is feasible and tolerated in unanesthetized rats and achieves clinically relevant buprenorphine exposure, enabling route-matched preclinical studies of buprenorphine pharmacology and opioid-related behavior.

## 1. INTRODUCTION

Opioid use disorder (OUD) remains a leading cause of preventable death in the United States (Garnett MF, 2026; Kaliamurthy et al., 2025). Buprenorphine (BUP), a partial mu-opioid agonist, is the most widely prescribed medication for OUD (Dahan et al., 2006; Kozinska et al., 2026). It is given as a sublingual film, usually combined with naloxone at 4:1 to discourage parenteral misuse, at a typical maintenance dose of 16 mg/day (Indivior Inc., 2025).

Preclinical rodent studies have delivered BUP subcutaneously (Chiang et al., 2010; Curtin et al., 2009), intraperitoneally (Wilson et al., 2016), intravenously (Yu et al., 2006), by osmotic minipump (Bossert et al., 2020; Tobacyk et al., 2023) or by oral gavage (Norden et al., 2024). None reproduces clinical sublingual delivery: parenteral routes bypass first-pass metabolism entirely, whereas gavage imposes the hepatic extraction that sublingual absorption is intended to avoid. Sublingual film administration has not, to our knowledge, been reported in rodents. The barriers are practical and anatomical because animals cannot be trained to retain a film, and the rodent oral mucosa is narrower and more keratinized than the human, which may limit absorption (Patel et al., 2011).

Here we report direct sublingual administration of the clinical BUP/naloxone film to unanesthetized rats. We characterized plasma BUP over 24 h at a dose scaled from the clinical maintenance regimen and assessed tolerability after 21 days of daily dosing, establishing the feasibility of route-matched BUP dosing in rats.

## 2. MATERIALS and METHODS

### 2.1 Animals

Male Sprague-Dawley rats (Charles River; 7-10 weeks) were group-housed under standard conditions with food and water *ad libitum*. Procedures were approved by the University of Pittsburgh IACUC (D16-00118; ID: 23124093).

### 2.2 Study design

Twenty rats received a single sublingual dose for PK analysis. A separate cohort (control n = 4, BUP n = 5) received 3.2 mg/kg BUP once daily for 21 days with daily body weights; terminal collection was on day 22, 24 h after the final dose. A target dose of 3.2 mg/kg BUP was derived from the 16 mg/day clinical maintenance regimen by the FDA body-surface-area conversion factor 6.2 (Nair and Jacob, 2016), adjusted for reported sublingual bioavailability in humans (∼30%)(Mendelson et al., 1997) and rats (∼13%) (Brewster et al., 1981). Clinical BUP/naloxone (2 mg/0.5 mg) films (Dr. Reddy’s Laboratories) were cut by scalpel to a size set daily from each rat’s body weight (mean 1.25 cm × (0.7 ± 0.4 cm)). Animals were restrained with a V-hold controlling the head and neck. Film pieces were placed and held sublingually with forceps for 60 s until softened, followed by brief scruffing to prevent expulsion or prematurely swallowing. The oral cavity was checked to confirm dissolution before release. Controls received an identical sham procedure without film.

### 2.3 Sample collection and measurements

To limit animal use without biasing estimates (Houston et al., 2021), 0–24 h profiles used staggered sparse sampling across three subgroups (n = 6–7): a warmed lateral tail vein draw at 1, 2 or 4 h paired with terminal cardiac puncture at 8, 12 or 24 h. Plasma BUP and norbuprenorphine (norBUP) were quantified by LC-MS/MS as previously described (Tobacyk et al., 2023), with terfenadine (10 ng/mL) as internal standard and an LLOQ of 0.02 ng/mL for both analytes; noncompartmental analysis used PKNCA (Denney, 2015). Terminal serum from repeat-dosed rats was assayed for AST, ALT and ALP (IDEXX BioAnalytics). Livers were removed intact at the hilum, blotted and weighed; the largest lobe was formalin-fixed and H&E-stained for modified Ishak scoring (Goodman, 2007).

### 2.4 Statistical analyses

Data are mean ± SD (PK) or mean ± SEM (all other measures). Body weight was compared by two-way ANOVA with Šidák *post hoc*; liver weight, liver-to-body-weight ratio, and serum enzymes by unpaired Welch’s t-tests. Significance was *p* < 0.05 (GraphPad Prism 11).

## 3. RESULTS

Plasma BUP peaked at 4.0 ± 3.4 ng/mL at 2 h and declined to 0.9 ± 0.3 ng/mL at 24 h (**Fig. 1**), all values exceeding the LLOQ by more than an order of magnitude. The peak fell within the 1.6–6.4 ng/mL reported at comparable times after sublingual dosing in humans (McAleer et al., 2003), and the 24 h concentration within the reported human range of 0.7–1.25 ng/mL (Bjornsson et al., 2023; Walsh et al., 2024). Dose-normalized AUC0-24was 12.2 h·ng/mL per mg/kg, within the 11.1–14.9 predicted for a 60–80 kg adult and within 10.0–20.6 when human sublingual bioavailability was varied across its reported SD (29 ± 10%) (Mendelson et al., 1997) (**Supplemental Table 1**). To test whether absorption occurred across the oral mucosa rather than in the gastrointestinal tract, we quantified norBUP, the major active metabolite (Elkader and Sproule, 2005), in the same samples. NorBUP remained well below parent drug throughout (**Fig. 1**), with AUC0-24of 5.4 versus 39.0 h·ng/mL and a metabolite-to-parent ratio of 0.14. Because norBUP is formed by hepatic N-dealkylation, this low fraction indicates that absorbed drug at least partly avoided first-pass metabolism.

**Figure 1.**
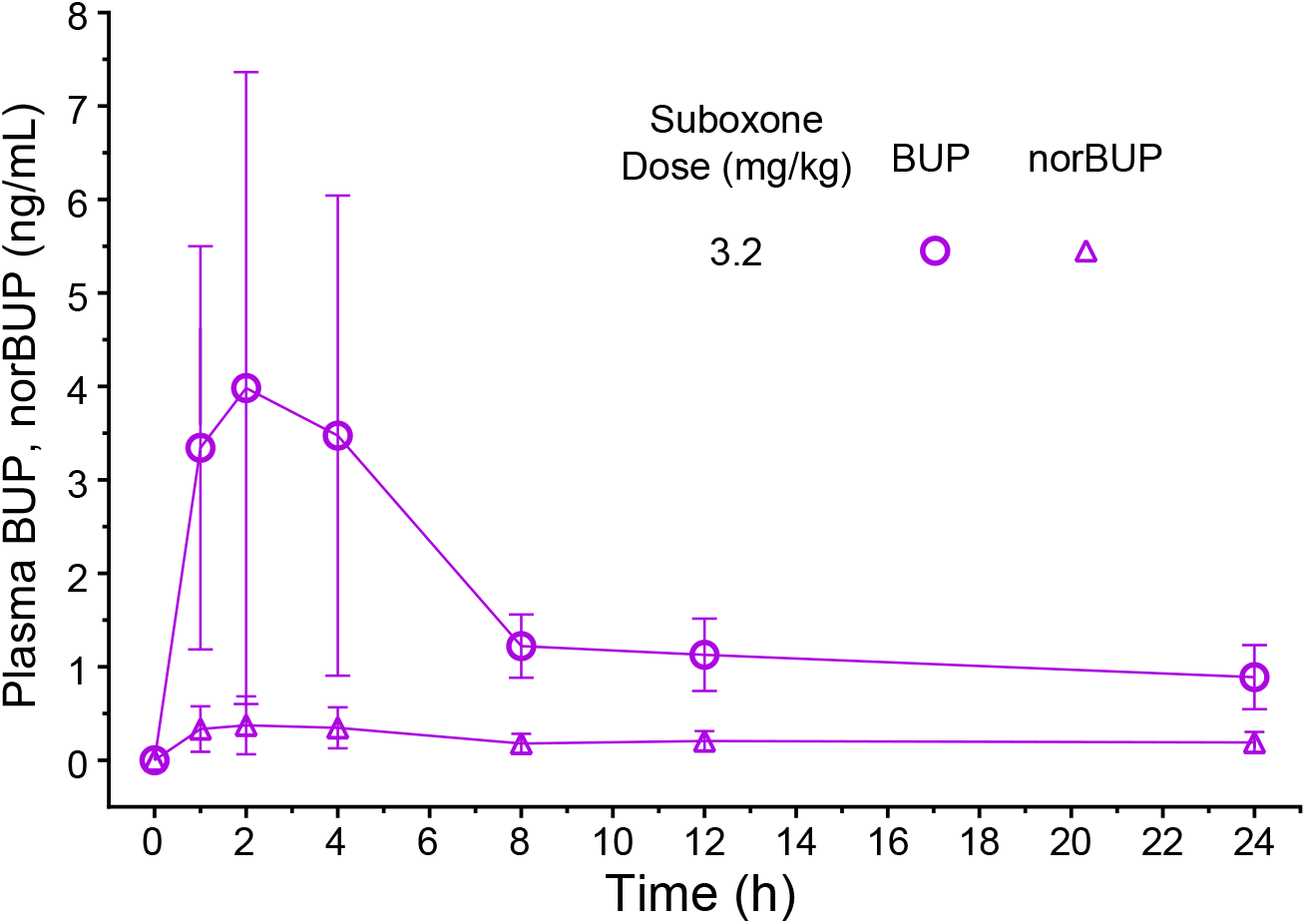
LC-MS/MS measurements of plasma buprenorphine and norbuprenorphine after sublingual administration of Suboxone. Plasma concentrations of BUP (magenta open circle) and norBUP (magenta open triangle) were sampled up to 24 h after a single BUP dose of 3.2 mg/kg (n= 6 or 7 per timepoint). Data are presented as mean ± SD.

Tolerability was assessed after once-daily dosing for 21 days. Both control (n=4) and BUP (n=5) groups gained weight, with a significant time × treatment interaction (repeated-measures two-way ANOVA, Geisser-Greenhouse corrected: F(2.088, 14.62) = 6.367, p = 0.0096; **Fig. 2a**). At terminal collection, treated rats showed an 8.4% lower body weight (242.8 ± 4.2 vs 265.2 ± 8.2 g; p = 0.063; **Fig. 2b**) alongside significant reductions in liver weight (7.70 ± 0.12 vs 9.35 ± 0.42 g; −17.6%; p = 0.025; **Fig. 2c**) and relative liver weight (3.17 ± 0.03 vs 3.52 ± 0.09%; −9.9%; p = 0.026; **Fig. 2d**). Welch’s test was used throughout because group variances differed.

**Figure 2.**
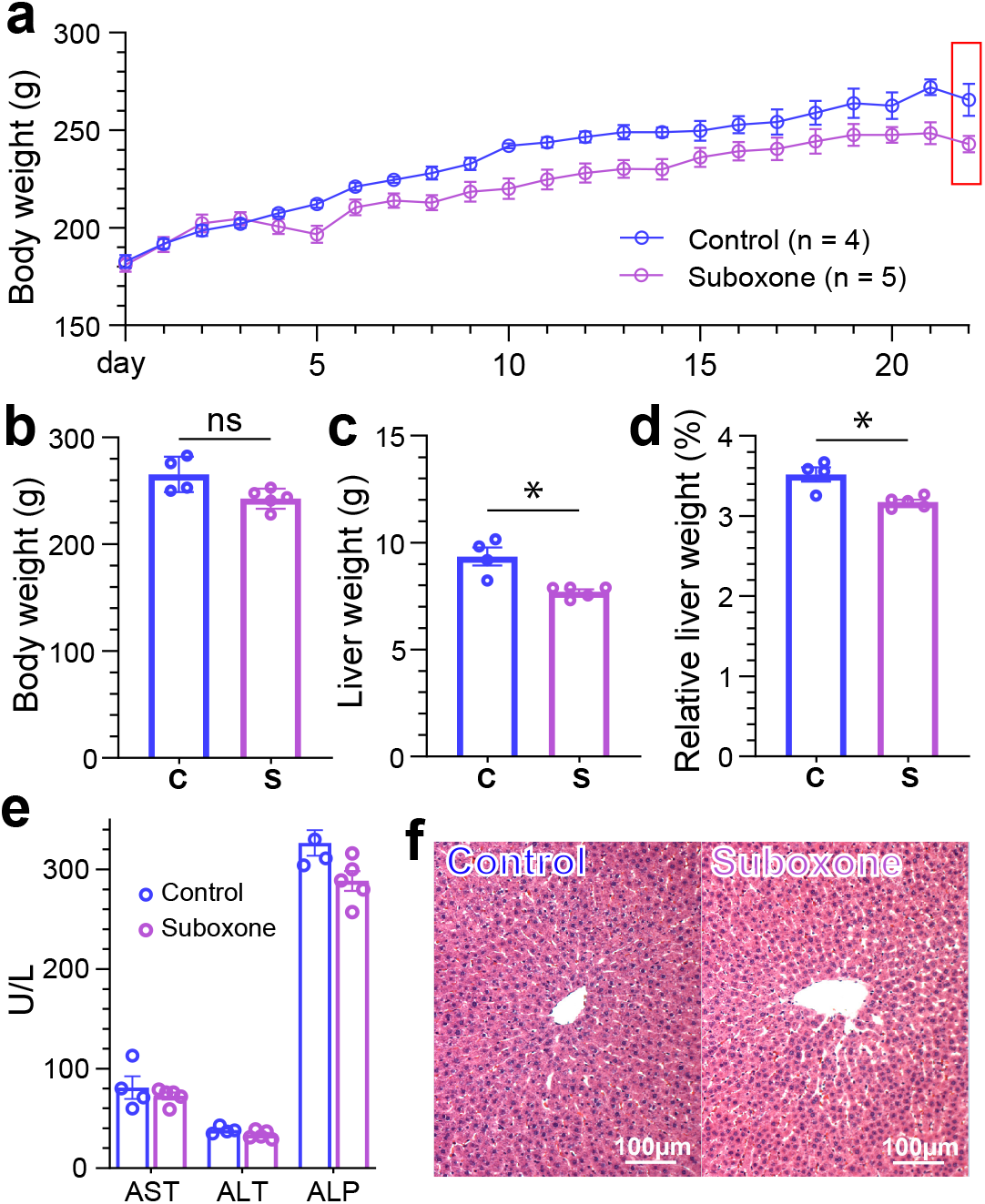
Body and liver tolerability to repeated sublingual Suboxone dosing. **(a)** Sham control or Suboxone of 3.2mg/kg (equivalent to clinical dose 16mg per day in humans) was sublingually administered daily from day 1 to day 21, showing a significant time × treatment interaction (repeated-measures two-way ANOVA, Geisser-Greenhouse corrected: F(2.088, 14.62) = 6.367, p = 0.0096). (**b**) Comparison of terminal body weights between sham control (C) and Suboxone-treated (S) rats (p = 0.063). **(c)** Comparison terminal liver weights (−17.6%; p = 0.025). **(d)** Comparison terminal relative liver weights (−9.9%; p = 0.026). **(e)** Repeated sublingual dosing over 3 weeks did not induce significant changes in activity of liver enzymes, including AST (p = 0.5162), ALT (p = 0.1100), and ALP (p = 0.0557). Welch’s *t* test was used throughout (b)-(e). Data are presented as mean ± SEM. **(f)** Representative H&E staining of liver tissues, showing no liver damage.

We also examined markers of hepatic injury because relative liver weight fell and the reduction in liver mass might not be attributable to lower body weight alone. Serum AST, ALT and ALP did not differ between groups (**Fig. 2e**). H&E sections scored by a blinded observer using the modified Ishak system (Goodman, 2007) returned 0 for every animal, with intact hepatocyte cords, normal sinusoidal architecture, and no inflammatory infiltrate, lipid accumulation or focal necrosis (**Fig. 2f**) (Thoolen et al., 2010). Therefore, liver mass declined without biochemical or histological evidence of hepatocellular injury or fibrosis.

## 4. DISCUSSION

Preclinical BUP studies have relied on routes that do not reproduce clinical sublingual delivery, and route governs both absorption kinetics and first-pass metabolism. We show that the clinical film can be administered sublingually to unanesthetized rats at a dose scaled from maintenance therapy, producing plasma BUP within the range reported in humans (McAleer et al., 2003) and, at 24 h, within the range associated with withdrawal suppression (Bjornsson et al., 2023; Walsh et al., 2024).

Rats did not retain a passively placed film, so the procedure depends on holding it beneath the tongue with forceps under gentle restraint for approximately one minute, by which point it reaches a semi-dissolved consistency, adheres to the mucosa, and is no longer expelled. Inter-animal variability was 23–85% across timepoints (Fig. 1), comparable to the 34–80% reported for sublingual BUP in humans (Mendelson et al., 1997; Santiago et al., 2020). Because each rat contributed a single measurement per timepoint, these values incorporate variability in the timing of peak concentration and may therefore overstate variability in total exposure.

Salivation raised the possibility that part of the dose was swallowed. The metabolite data argue against a major enteral contribution: norBUP accounted for only 14% of parent exposure, consistent with absorption that partially bypasses hepatic first-pass metabolism. Glucuronide conjugates were not quantified and would be needed to assess total first-pass metabolism. Two further limitations bear on the PK data. Early samples came from the tail vein in awake rats and later samples by cardiac puncture under isoflurane, so sampling route and anesthetic exposure covary with time — relevant because isoflurane reduces hepatic blood flow for a high-extraction drug.

Sampling to 24 h under a sparse design also precluded characterization of the terminal phase.

Twenty-one days of dosing produced no biochemical or histological evidence of hepatic injury, but liver weight fell 17.6% and relative liver weight 9.9%. The most likely explanation is nutritional and metabolic adaptation rather than toxicity (Mezencev et al., 2024). Opioids including BUP suppress appetite (Jirkof et al., 2015; Wiffen et al., 2014), and reduced caloric intake depletes hepatic glycogen and lipid stores without cellular damage; reduced water intake may also lower the mass of this highly vascularized organ (Diaz-Munoz et al., 2010), and lower hepatic metabolic demand can produce reversible hepatocyte atrophy rather than cell loss (Belloni et al., 1988). Food and water intake were not measured here, which is the clearest test of this interpretation and should be included in future work. Within these limits, sublingual film dosing is feasible and tolerated in unanesthetized rats and achieves clinically relevant BUP exposure, enabling route-matched preclinical studies of BUP pharmacology and opioid-related behavior.

## CRediT authorship contribution statement

Yao Wang: Investigation, Methodology, Formal analysis, Visualization, Data curation, Writing – original draft, review and editing. Qiang Chen: Investigation, Methodology, Formal analysis, Visualization, Data curation, Writing – review and editing. Thomas Deakin: Investigation, Methodology, Formal analysis, Visualization, Data curation, Writing – review and editing. Trent Emerick: Methodology, Conceptualization, Writing – review and editing. Yan Xu: Writing – Methodology, Conceptualization, Writing – review and editing. Funding acquisition, Formal analysis. Pei Tang: Conceptualization, Funding acquisition, Project administration, Supervision, Methodology, Visualization, Formal analysis, Writing – original draft, editing and review.

## Funding sources

The reported research was supported in part by the US National Institutes of Health (NIH, R21DA062076-01A1 to P.T. and R01NS122830 to Y.X.). The NIH had no role in the study design, data collection and interpretation, or the decision to submit the work for publication.

## Declaration of competing interest

The authors declare that they have no known competing financial interests or personal relationships that could have appeared to influence the work reported in this paper.

## Acknowledgment

The authors are grateful to Dr. Jacques Chelly of University of Pittsburgh School of Medicine for his valuable input at the early phase of the project.

## Appendix Supplementary materials (1)

Table 1

## Notes

### Competing Interest Statement

The authors have declared no competing interest.

