## Supplemental Table 1 for "Sublingual Buprenorphine/Naloxone Film in Unanesthetized Rats: Feasibility, Tolerability, and Plasma Exposure"

**Supporting Table 1.** Derivation of the rat sublingual dose and predicted systemic exposure from published human data.

**A. Input parameters**

| Parameter | Value | Source |
|---|---|---|
| Clinical maintenance dose, buprenorphine | 16 mg/day | (Harris et al., 2026) |
| Human reference body weight | 70 kg (range 60–80) | Assumed |
| Human dose-normalized plasma buprenorphine $AUC_{0-72}$, 16 mg sublingual | 2.66 h·ng/mL per mg | (Gu et al., 2023) |
| Body-surface-area dose conversion factor, human → rat | 6.2 | (Nair and Jacob, 2016) |
| Sublingual bioavailability, human | 30% (29 ± 10%) | (Mendelson et al., 1997) |
| Sublingual bioavailability, rat | 13% | (Brewster et al., 1981) |
| Sublingual bioavailability ratio (human/ rat) | 2.31 | Calculated |

**B. Dose derivation**

| Step | Operation | Result |
|---|---|---|
| 1 | Clinical dose ÷ body weight = 16 ÷ 70 | 0.229 mg/kg |
| 2 | × BSA factor 6.2 | 1.42 mg/kg |
| 3 | × bioavailability ratio 2.31 | 3.27 mg/kg |
| 4 | Rounded to administered dose | **3.2 mg/kg** |

**C. Predicted rat exposure**

| Step | Operation | Result (h·ng/mL per mg/kg) |
|---|---|---|
| 1 | Human $AUC_{0-72}$ per mg × 70 kg = 2.66 × 70 | 186.2 |
| 2 | ÷ BSA factor 6.2 | 30.0 |
| 3 | ÷ bioavailability ratio 2.31 | **13.0** |

**D. Sensitivity of the prediction**

| Parameter varied | Range tested | Predicted exposure |
|---|---|---|
| Human body weight | 60–80 kg | 11.2–14.9 |
| Human sublingual bioavailability | 19–39% (mean ± 1 SD) | 10.0–20.6 |
| Rat sublingual bioavailability | 13% with no reported variance | |

**E. Observed versus predicted**

| Quantity | Value |
|---|---|
| Observed $AUC_{0-24}$, 3.2 mg/kg | 39.0 h·ng/mL |
| Observed, dose-normalized | 12.2 h·ng/mL per mg/kg |
| Predicted at reference parameters | 13.0 |
| Ratio, observed / predicted | 0.94 |
